# Extended normothermic machine perfusion preserves viability and tissue integrity of ex vivo human intestine segments

**DOI:** 10.64898/2026.09.17.752436

**Authors:** Pamela A. Manco Urbina, Sae Rome Choi, Dillon C. Cheung, Lena K. Egbert, Marlene I. Garcia-Neuer, Stephanie Yu, Nisha S. Rehman, Danielle Hanks, Tuan-Anh Le, Husnu H. Alabay, Kelsey E. McHugh, Feda H. Hamdan, Nimesh Naik, William A. Faubion, Justin T. Brady, Hakan Ceylan, Suman Bose

## Abstract

Human intestinal diseases, including inflammatory bowel disease (IBD), are difficult to model because existing animal and in vitro systems do not capture the intestine’s integrated vascular, immune, and metabolic complexity. Here, we establish a custom normothermic machine perfusion platform that maintains surgically resected human intestinal tissue for up to 72 hours under near-physiological conditions. Discarded specimens from patients with nonmalignant intestinal diseases, including IBD and diverticulitis, were continuously perfused at 37°C with real-time hemodynamic and metabolic monitoring. Perfusion preserved tissue viability, mucosal architecture, epithelial integrity, oxygen consumption, metabolic activity, and vascular patency. Daily perfusate sampling enabled longitudinal profiling and experimental modulation of inflammatory mediators, revealing patient-specific immune states. This long-duration platform bridges reductionist in vitro models and clinical disease, enabling mechanistic studies of intestinal inflammation and future testing of personalized therapies in intact patient-derived tissue.

## INTRODUCTION

Intestinal diseases such as inflammatory bowel disease (IBD) impose a major global health burden, affecting hundreds of millions of individuals and contributing substantially to chronic morbidity, healthcare utilization and reduced quality of life.^1–4^ These disorders arise from complex and dynamic interactions among epithelial barrier dysfunction, dysregulated immune responses, altered metabolism, and microbial microenvironments.^5–10^ Intensive research efforts have yielded a diverse range of therapeutics, including antibodies, small-molecule inhibitors, immune modulators, and experimental cell therapies. Yet patient responses remain variable as roughly only 20 - 40% go into clinical remission,^11–13^ and many who initially respond subsequently lose response. This therapeutic ceiling drives treatment failure, leaving a substantial fraction of patients dependent on surgical intervention. A major barrier to development of personalized precision medicine in IBD is the lack of preclinical drug testing platforms that faithfully recapitulate the integrated physiology of the human intestine.

Current preclinical models of IBD have a limited ability to capture the complex immunological, genetic, and environmental features of human gastrointestinal tissue. In vivo models have been the most widely used preclinical platform for testing IBD therapeutics. Early models of experimental colitis relied on chemical agents to reproduce the symptoms of the disease.^14–17^ More recent efforts have focused on recapitulating specific immunological pathways, either through gene knockouts such as IL-10 or knock-ins that mimic human polymorphic variants underlying defective epithelial barrier function^18–20^. Humanized mouse models bring greater relevance to the human immune system and have proven effective for testing immunomodulatory therapeutics.^21,22^ However, even in these models, human immune cells fail to interact properly with rodent stromal components, and these models lack the microbial components central to IBD, limiting faithful replication of human pathobiology. As a result, drugs that often succeed in animal models often fail to predict patient outcomes or the variability seen across patient populations.

In vitro microphysiological systems have recently emerged to bridge the gap between rodent models and human disease. Human colon crypt-derived organoids have served as a foundational platform for studying epithelial biology and for drug testing,^23,24^ and recent advances in stem cell engineering have enabled multicomponent systems comprising epithelial, neural, and vascular lineages derived from gut stem cells.^25^ A major limitation of organoids, however, is the absence of human immune components, which precludes their use as faithful disease models. Organ-on-chip systems, which employ cell lines and can incorporate immune cells, have not yet reached the sophistication needed to reproduce the complex multicellular interactions that define a disease context.

Intact whole-bowel segments represent an ideal test bed for replicating gastrointestinal biology and an ideal platform for drug testing. Ex vivo static culture of whole swine intestinal segments has been used to predict the absorption of orally administered drugs^26^ and a rodent intestinal organ culture system enabled the first demonstration that microbes induce regulatory T cells.^27^ Whole-tissue segments obtained directly from patient samples incorporate all the essential biological components relevant to the disease - the epithelial barrier, an autoreactive immune system, stromal cells, vasculature for cell trafficking, and the patient’s own diverse microbiota. As such, they would constitute the closest model to a human clinical trial and could serve as a final test bed for drug efficacy. Maintaining large pieces of human tissue is challenging, however, both surgically and in terms of preserving viability.

Normothermic machine perfusion (NMP) has emerged as a transformative approach for maintaining human organs *ex vivo* under near-physiological conditions. NMP has revolutionized solid organ transplantation by enabling preservation of liver,^28–32^ kidney,^33,34^ heart,^35^ and lung^36^ for several days. The ability to sustain organ-level physiology ex vivo using NMP creates a unique platform to evaluate therapeutics under near-physiological conditions prior to test in human.^37–39^ However, despite these advancements, the application of NMP to the human intestine remains strikingly underdeveloped. Early studies focused on short-term preservation of canine intestinal segment for transplantation^40,41^ and human intestinal for absorption mechanistic studies.^42^ In both cases the technical feasibility of intestinal perfusion was demonstrated, but lacked systematic evaluation of tissue viability, immune preservation, or long-term metabolic function. More recent studies have focused primarily on porcine intestine, but have been restricted to short perfusion durations of less than 6 hours and focused on trasplantation^43^, which is not optimal for therapeutic testing. Hypothermic perfusion approaches have also been explored for intestinal preservation^44^, however, these systems do not support in vivo-like metabolism or immune activity and are therefore poorly suited for modeling inflammatory intestinal diseases.

Here, we report the feasibility of a custom normothermic machine perfusion platform capable of sustaining human intestinal specimen ex vivo for 48-72 hours under controlled, normothermic conditions. Using human small and large bowel specimens obtained from patients undergoing resection for benign intestinal diseases, including IBD and diverticulitis, we successfully cannulated native mesenteric vasculature and established stable perfusion at 37°C with continuous monitoring of pressure, flow, oxygenation, and metabolic parameters. By enabling prolonged maintenance of human intestinal tissue with preserved immune and vascular compartments, this platform provides a foundation for mechanistic interrogation of intestinal inflammation and a translational testbed for evaluating therapeutics in an intact patient-derived context. Together, these findings establish the feasibility of extended normothermic perfusion of the human intestine for future applications in personalized disease modeling and preclinical evaluation of targeted interventions.

## RESULTS

### Human Intestinal Specimens and Experimental Cohort

Human intestinal specimens were obtained from patients undergoing segmental colectomy or small bowel resection for benign disease, including diverticulitis and inflammatory bowel disease (IBD). While inter-sample variability was observed across the specimens, this variability reflected intrinsic biological and microbiological differences rather than technical failure of the perfusion platform. In total, 17 specimens comprising colonic and small bowel tissues were evaluated. Eleven specimens were used during protocol development and optimization (data not shown), resulting in the finalized NMP protocol implemented in this study. Six specimens were subsequently perfused using the finalized NMP configuration, with all six completing the experimental protocol without technical failure.

For the assessment of metabolic activity, data from the six specimens are presented as grouped longitudinal trends to capture the collective performance of the platform. To provide a high-resolution view of the histological integrity and divergent immune responses, two specific cases—one diverticulitis-derived colon and one IBD-derived small bowel—were selected as representative phenotypic benchmarks. Comprehensive data for all additional individual cases are provided in the Supplementary Information.

### Tissue harvest and normothermic machine perfusion (NMP) setup

Our normothermic machine perfusion (NMP) system was established to enable continuous delivery of oxygen and nutrients to the human intestinal specimens under controlled, near-physiological conditions. The organ chamber and perfusate reservoir were maintained within a humidified incubator maintained at 37 °C, providing stable normothermic conditions with ambient air oxygen levels and 5% CO_2_ throughout the perfusion period. A schematic of the perfusion setup is shown in **Figure 1A**, consisting of a peristaltic pump driving perfusate flow and inline sensors enabling real-time monitoring of oxygen tension, flow rate, and pressure at the inlet. Gas permeable bags were used as media reservoir to enable rapid gas exchange of the perfusate. Real-time monitoring of perfusion parameters enabled timely adjustments, particularly modulation of flow in response to increases in pressure.

**Figure 1.**
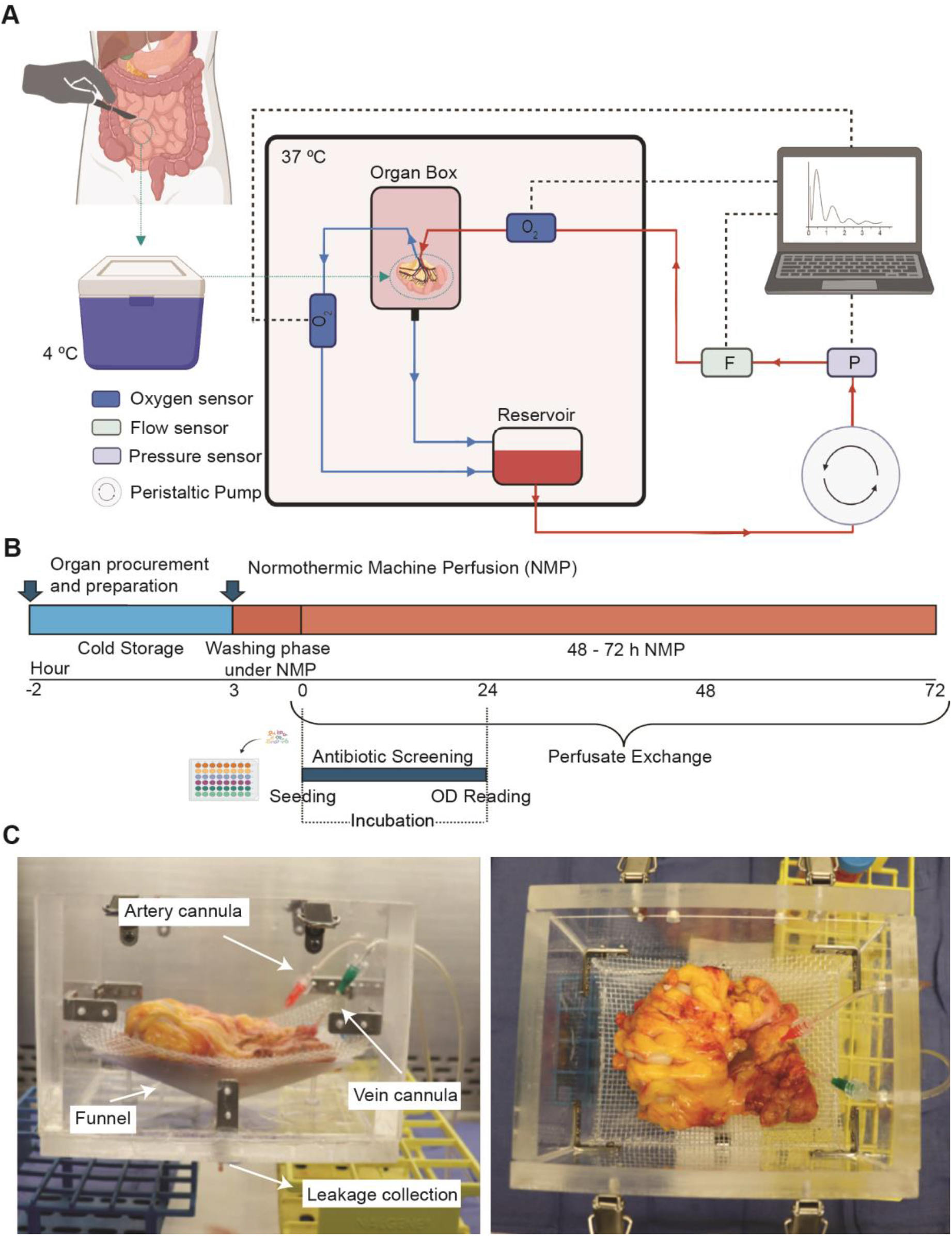
Setup and experimental design. **(A)** Schematic of the experimental outline, starting from specimen procurement, subsequent transportation under cold conditions, and final normothermic machine perfusion with real-time monitoring of several perfusion parameters. **(B)** Workflow of the overall experimental design, first 2 hours of cold storage, encompassing, specimen procurement, transportation and preparation, followed by 48 to 72 hours of normothermic perfusion (NMP); in parallel, an infection control screening testing several antibiotics for a personalized perfusion system. **(C)** Side view of organ box **(D)** Top view of the organ box, displaying the organ funnel and mesh.

The overall timeline is summarized in **Figure 1B**. Patients undergoing colon or small bowel resection for benign diseases including Crohn’s Disease, Ulcerative Colitis or diverticulitis were recruited for participation from the colorectal surgeons’ practices. Patients received a mechanical bowel preparation with oral antibiotics (neomycin and flagyl) per the standard of care preoperatively. The patients then underwent resection of the bowel per the standard of care for the operating surgeons. Operations were performed in an open or minimally invasive fashion at the surgeon’s discretion. The extent of bowel resection was not altered for the study. Resections were performed with a high ligation of the feeding mesenteric vessel (inferior mesenteric artery, ileocolic artery, proximal small bowel arcade). After resection, the bowel was transferred to a sterile draped table and the intestine specimen partially opened longitudinally. The specimen was inspected by the study team, surgeon and board-certified pathologist to confirm that a segment of bowel did not require microscopic inspection. The planned segment to be used in the study was approximately 3-4 cm in length and left circumferentially intact and everted for thorough inspection. The feeding artery and vein were identified in the mesentery close to the root of the feeding vessel. These were isolated and cannulated separately using 14 - 24 gauge angiocatheters based on vessel size and secured with 3-0 silk ties. The mesentery was then divided using a handheld Ligasure^TM^ (Medtronic, Minneapolis, MN, USA) from adjacent to the cannulated vessels to the bowel wall at the previously defined borders of the chosen specimen. The bowel was then divided sharply with scissors. The cannulated artery was flushed to confirm veinous return and identify any areas of leak from the cut edge of mesentery. Areas of leak were controlled with the Ligasure or clamps and ties.

The specimen was transported in Belzer cold storage solution supplemented with penicillin/streptomycin on wet ice. Typical cold ischemia time from removal of the segment of bowel from the study participant to being placed on pump was 2 –3 hours, depending on the specimen complexity. Key challenges in the delay resulted during the cannulation of the specimen, due to small vessel sizes often seen with small bowel-derived specimens and the presence of large amounts of adipose tissue within the mesentery for sigmoidal colon-derived specimens, which complicated vessel identification and increased preparation time. Upon arrival in laboratory, the tissue was first imaged using fluoroscopic angiography to map perfusion path (see next section). Next, it was washed several times with cold saline to remove any remaining stool and tissue debris and placed in the NMP circuit. The NMP process started with a washing phase, where the specimen was kept in an organ bag and immersed in an acellular perfusate supplemented with antibiotics and antifungals in an open circuit. We found that this washing step dramatically decreased chances of infection of the sample. Following the washing phase, the tissue was moved to a custom organ box to support tissue positioning and effluent collection. The organ box was designed together with a 3D-printed funnel to collect passive leakages (**Figure 1C**). The intestinal specimen was placed on a nylon mesh to prevent cannula obstruction and minimize mechanical stress on the tissue (**Figure 1D**). NMP was initiated by adding fresh perfusate to reservoir and enabling recirculation of perfusate from the tissue outlet back to the reservoir bag. The perfusate consisted of IMDM cell culture media supplemented with fetal bovine serum (FBS) and N-acetyl L-cysteine and valproic acid, to mitigate oxidative stress and inflammatory responses, respectively.^45–49^

The room temperature module consisted of a peristaltic pump driving the perfusate from the reservoir to the intestinal specimen, followed by in-line flow and pressure sensors to monitor perfusion parameters. A bubble trap was positioned downstream from the pressure sensor to remove any air bubbles prior to the tissue arterial cannula. Within the incubator, the perfusion circuit incorporated an in-line oxygen sensor proximal to the arterial cannula, while tissue effluent was collected through the custom funnel inside the organ box, and returned to the reservoir in a closed-loop circulation. A water bath was placed inside the incubator to prewarm the perfusate to 37 °C immediately before entering the tissue, ensuring normothermic perfusion.

### Evaluating tissue perfusion pre and post NMP

Fluoroscopic angiography is a safe and non-invasive procedure and routinely used in clinical practice to map vasculature through deep tissue and has recently been incorporated into *ex vivo* organ machine perfusion to assess vascular integrity and perfusion.^50,51^ Accordingly, we decided to use this technique to image the perfusate distribution through our sample and evaluate changes in tissue perfusion over the course of NMP. Specifically, we wanted to ensure that the cannulated vessels could perfuse the entire vascular network and detect and repair any major leaks in the perfusion path. Fluoroscopic angiography was performed on hypothermic samples before loading them on NMP system. Samples were flushed with undiluted Optipaque^TM^ iodinated contrast agent and imaged using an OEC 3D C-arm imaging system. Details of the procedure can be found in methods section. Representative angiography from an IBD-derived specimen demonstrated widespread perfusion of the native mesenteric vasculature, reaching the fine capillary networks throughout the intestinal tissue (**Figure 2C**). Minor vascular extravasation was observed at the distal mesenteric margins, most likely due to minor tissue transections during surgery (**Figure 2C**, top). The sites of vascular effluence identified from the fluoroscopic angiographies were closed using a surgical tissue sealant (Preveleak®) to minimize primary focal leakages and ensure stable vasculature perfusion. After 48 hours of NMP, repeat angiography was performed to evaluate any changes in vascular perfusion that occurred during the experiment. This revealed a more heterogeneous perfusion compared with pre-perfusion images, as characterized by the attenuated contrast filling in central regions of the specimen (**Figure 2C**, bottom). Additionally, increased accumulation of vascular extravasation sites was observed close to the intestinal tissue.

**Figure 2.**
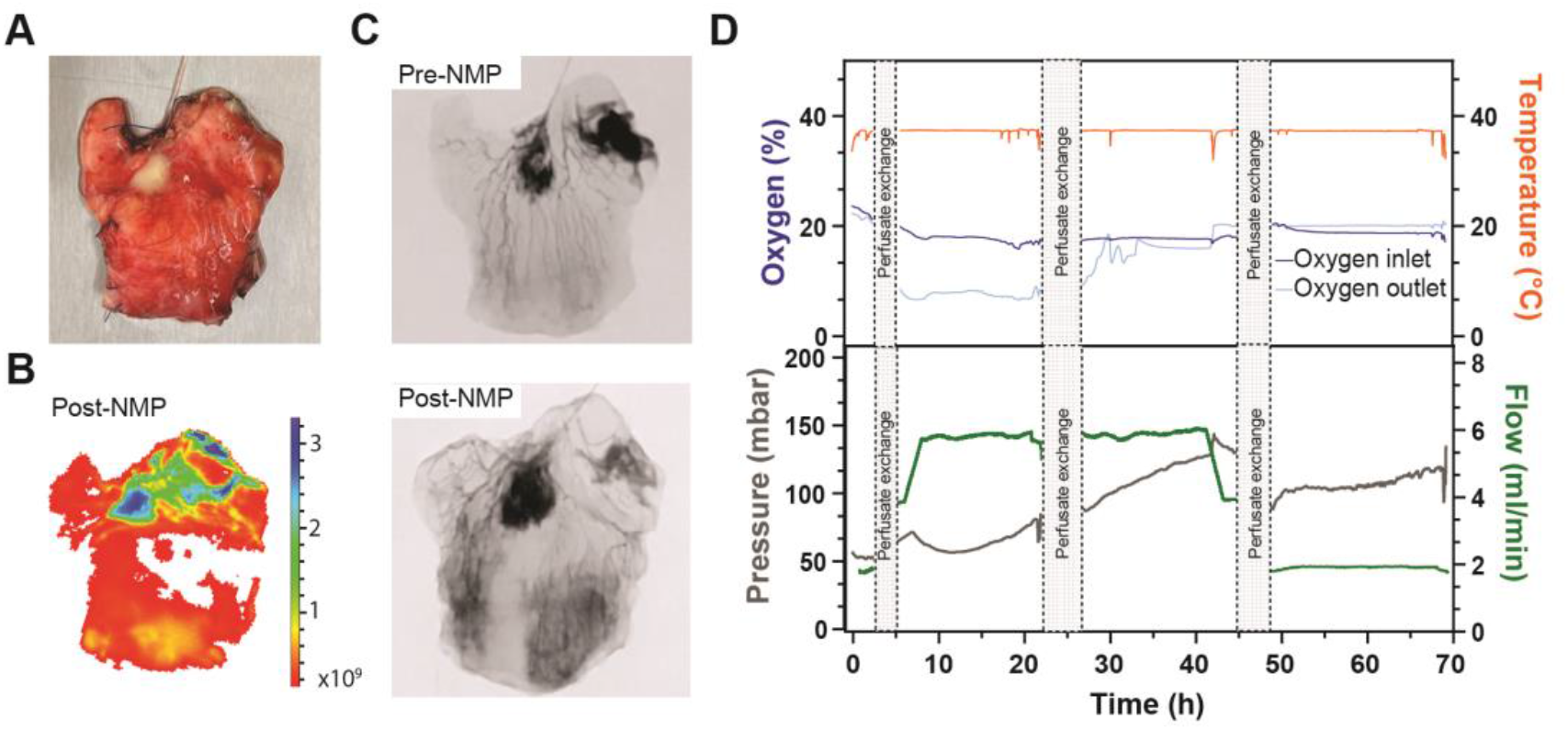
Perfusion machine performance. **(A)** Picture of the small bowel specimen before perfusion, showing arterial cannulation. **(B)** Representative near-infrared fluorescence image of the perfused specimen after 48 hours of perfusion, captured 30 minutes post-injection of indocyanine green (ICG). **(C)** Representative angiogram of the native mesenteric and intramural vasculature. **(D)** Representative longitudinal monitoring of perfusion parameters, including oxygen levels, temperature, flow, and pressure. All parameters were standardized to support tissue viability.

IVIS imaging system was used as a complementary technique to evaluate whole specimen perfusate distribution and microvascular perfusion beyond the spatial resolution of fluoroscopic angiography. At the end of NMP, indocyanine green (ICG) was perfused through the specimen’s native vasculature in combination with bovine serum albumin (BSA). The fluorescent dye binds to BSA, being predominantly confined in the intravascular space, enabling visualization of vascular perfusion throughout the tissue, including the microvascular network.52 A representative image of this technique is presented in Figure 2B, showing homogeneous perfusion across most of the tissue surface. Notably, increased fluorescence intensity was observed in regions that overlapped with those of vascular effluence identified from angiography (**Figure 2B-2C**). Across the full cohort, fluoroscopic angiography and IVIS imaging consistently demonstrated successful vascular perfusion both before and after NMP.

The custom perfusion platform maintained homeostatic conditions throughout the 48 –72 hour duration, named 48 h Protocol and 72 h Protocol respectively, as evidenced by stable temperature, consistent oxygen levels and controlled pressure, which were regulated to ensure optimal tissue viability. Perfusate flow rate was manually adjusted in response to changes in oxygen tension and arterial pressure to maximize oxygen delivery while minimizing the risk of tissue hypertension. Typical starting flow rate across all samples was 4 mL/min, which was adjusted to between 2 and 6 mL/min to address oxygen and pressure changes (Table S1). In this representative example displayed in Figure 2D, oxygen tension, measured at the artery inflow line, remained stable at approximately 18% throughout the perfusion period. In contrast, the oxygen levels measured at the venous outflow line showed a slow decrease for the first 5 hours, stabilizing at ∼7% pO2, indicating sustained oxygen consumption during perfusion. Oxygen extraction calculation was feasible only for this specimen, in which a stable venous return was achieved, allowing simultaneous measurement of arterial and venous oxygen saturation during the first 24 hours of perfusion. Beyond this time point, consistent venous return was interrupted, as a consequence, perfusate at the venous outlet equilibrated with ambient oxygen, showing similar values as inlet oxygen. Therefore, further calculation of oxygen consumption was not possible. Despite this limitation, the available data demonstrated sustained metabolic demand, with consumption of 0.028 mL Oxygen/mL/g bowel oxygen. Arterial oxygen saturation subsequently equilibrated to baseline levels, facilitated by passive oxygen diffusion into the perfusate through the gas-permeable reservoir bag and tubing. Moreover, over a 40-hour period, a continuous increase in arterial pressure was observed and thus the flow rate was reduced to a final rate of 2 mL/min to manage the pressure; this was critical to preserve vascular integrity while ensuring continuous oxygen and nutrient delivery.

We noted that by the end of perfusion about half of samples exhibited an approximately 40% increase in tissue weight, consistent with interstitial edema and highlighting the sensitivity of intestinal tissue to perfusion pressure dynamics over extended time.

### Personalized infection control for bowel samples

Infection control is a critical determinant of successful NMP and preservation of tissue viability. Human intestinal specimens are particularly challenging because microbial contamination may arise during surgery, cannulation, and sample transport, while each patient’s native intestinal microbiota and prior antibiotic exposure introduce substantial variability in microbial burden and antimicrobial susceptibility. In our initial experiments, specimens treated uniformly with penicillin-streptomycin and ciprofloxacin showed highly variable outcomes. Once overt infection developed, tissue viability did not recover despite additional antibiotic dosing. To better control the endogenous microbial load during prolonged NMP, we implemented a dynamic, specimen-specific antimicrobial screening protocol performed in parallel with perfusion. A schematics of the infection control strategy is depicted in **Figure 3A**, briefly, perfusate collected after the initial washout phase was screened against a panel of 11 individual antibiotics and defined antibiotic combinations, comprising 66 total antimicrobial conditions, using a 48-well plate format. Bacterial growth was assessed at 0, 24, and 48 hours by measuring optical density at 600 nm, enabling selection of the most effective antimicrobial regimen for each individual intestinal specimen. The screening data at the 24-hour mark provided a clear map of bacterial susceptibility and resistance. As shown in the absorbance heatmaps in **Figure 3B**, the most robust infection control was achieved through synergistic combinations. Specifically, the pairing of Piperacillin/Tazobactam with Penicillin/Streptomycin (Pen-Strep) frequently resulted in the lowest OD_600_ values, indicating a substantial arrest of metabolic activity from contaminating flora.

**Figure 3.**
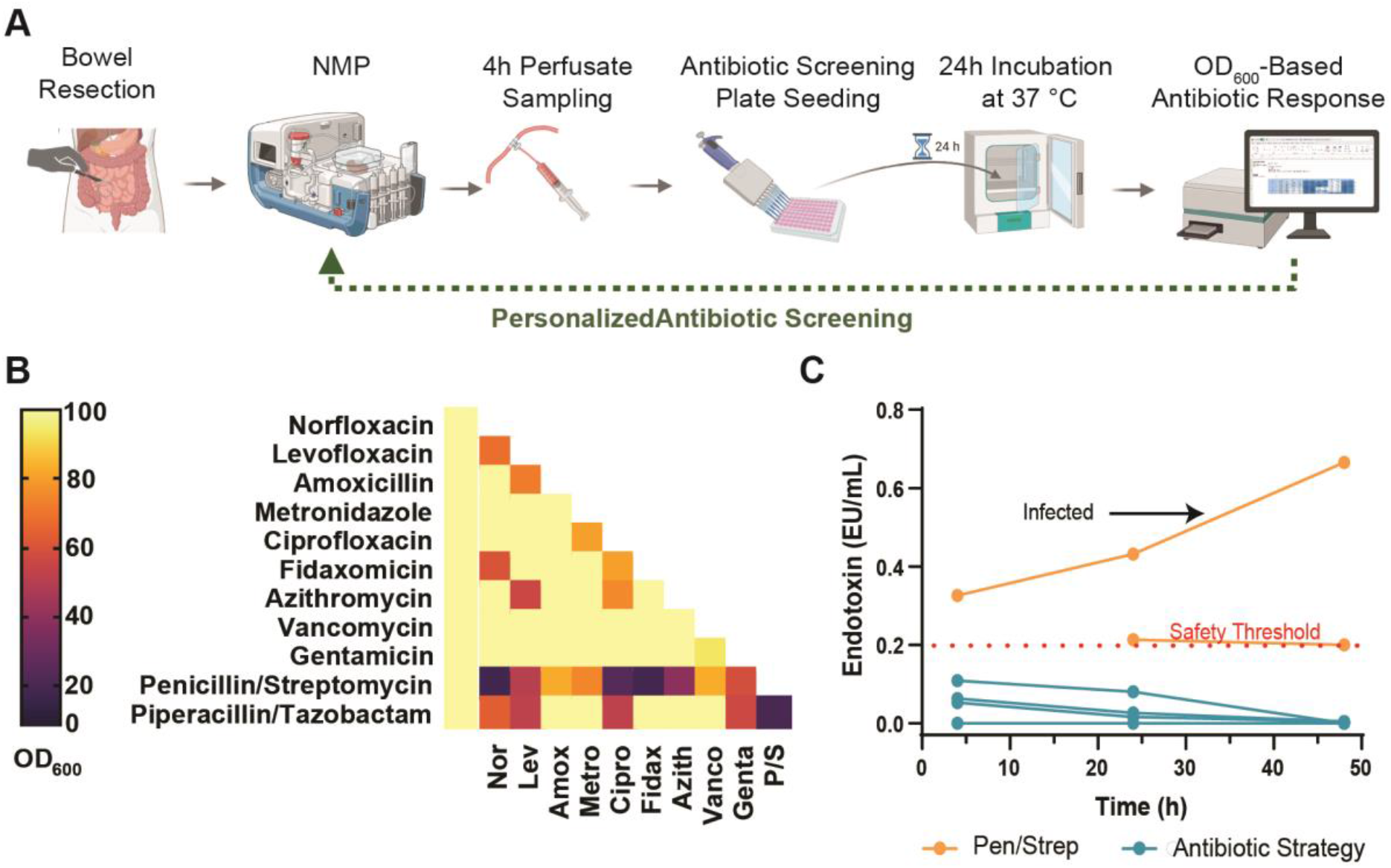
Antibiotic screening and infection assessment. **(A)** Graphical description of the Infection Control Strategy. **(B)** Representative heatmap of single and synergistic antibiotic combinations, displaying optical density (OD600) after 24 hours of incubation with “infected” perfusate. Data were normalized to baseline (T0) and are expressed as percentages. **(C)** Longitudinal profile of bacterial byproduct, Endotoxin, of 6 independent experiments, categorized by implementation of Infection Control strategy. Data is plotted by time point.

In instances where the initial microbial load was insufficient to reach detectable thresholds within 20 hours, a retrospective screening was performed to validate the chosen combination. Frozen perfusate from the washing step was used to perform the antibiotic screening, by waking up bacteria for 24 hours, and using the resulting solution for seeding the antibiotic plates and proceeding as before mentioned.

Systemic bacterial load was quantified by measuring endotoxin in perfusate. Optimized antibiotic treatment progressively led to complete control of infection in 4 out of 4 cases where it was implemented while two cases where Pen-Strep was used led to variable outcomes with one sample having uncontrolled bacterial contamination by the end of the study (**Figure 3C**). This targeted intervention ensured that the subsequent longitudinal secretome analysis remained reflective of endogenous tissue signaling rather than microbial interference.

### Evaluation of metabolic activity and cell injury

Glucose consumption, lactate production, and lactate dehydrogenase (LDH) release were selected as key perfusion biomarkers to assess organ metabolic activity and cell injury based on prior studies in solid organ normothermic perfusion.^28,52–54^

As shown in **Figure 4**, both glucose consumption rate (mg glucose/hour/gram of tissue) and lactate production (mmol lactate/g tissue) increased during the first 24 hours of NMP, followed by stabilization between 24 and 48 hours. Metabolic biomarkers are reported per time point rather than cumulatively. Given the intrinsic variability among specimens, including differences in metabolic activity and microbiota, the glucose consumption rate measured after the first 4 hours of perfusion was defined as the basal glucose consumption rate.

**Figure 4.**
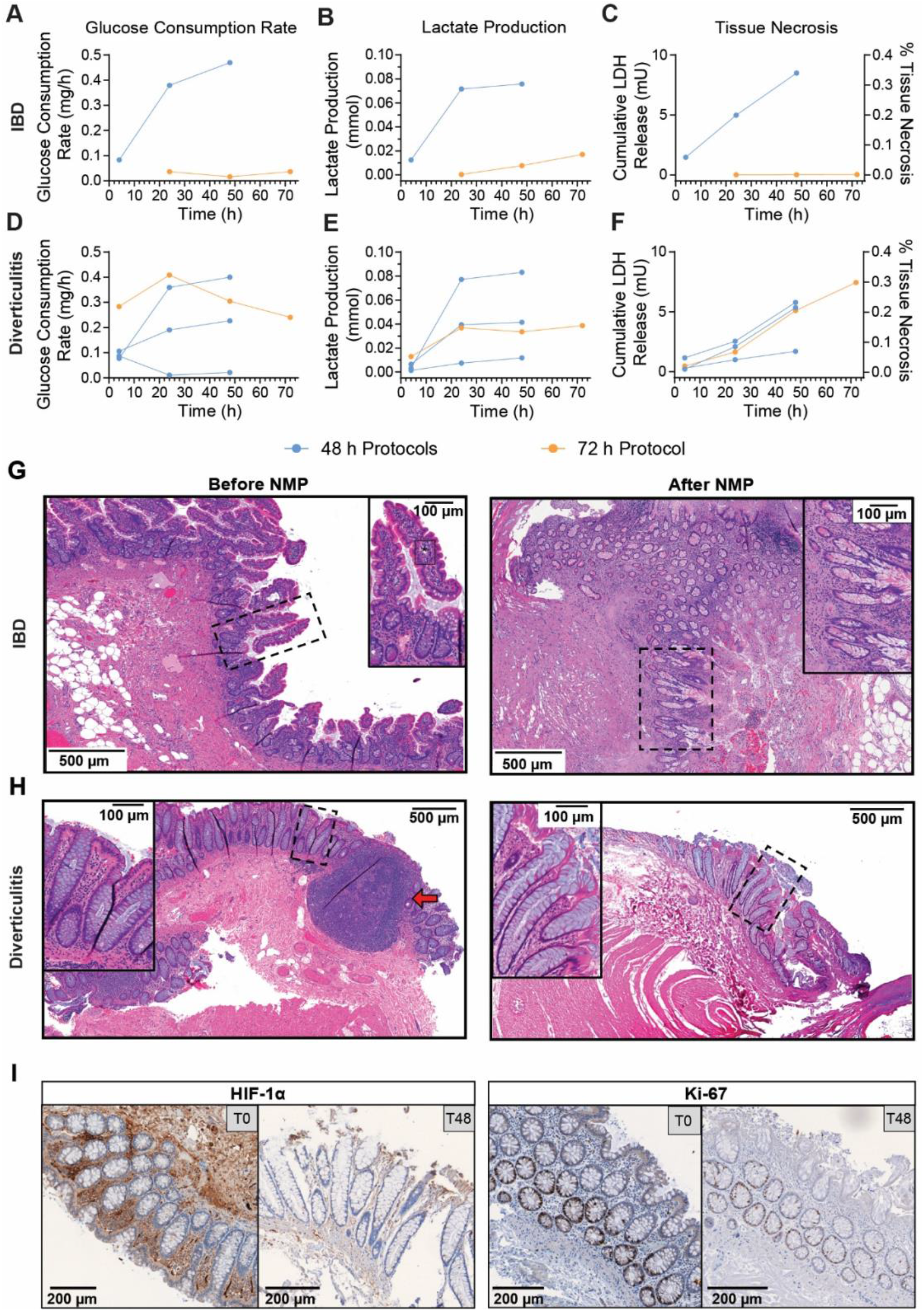
Metabolic stability and histological integrity of human intestinal specimens during NMP. **(A–F)** Longitudinal assessment of metabolic and cellular injury markers across the experimental cohort (n=6), with all values normalized to tissue weight (g). Glucose consumption rate (mg/h/g) of A) IBD-derive specimen and D) diverticulitis-derived specimen, lactate production (mmol/g) are presented per individual time point (4h, 24h, 48h), showing sustained metabolic activity of B) IBD-derive specimen and E) diverticulitis-derived specimen. Cumulative LDH release (U/g) over the 48 and 74-hour perfusion period as a marker of cellular viability and its conversion to percentage of tissue necrosis from C) IBD-derive specimen and F) diverticulitis-derived specimen. **(G)** Representative H&E-stained sections of an IBD-derived specimen before perfusion (left) and after 48 hours of NMP (right). **(H)** Representative H&E-stained sections of a diverticulitis-derived specimen before perfusion (left) and after 48 hours of NMP (right), demonstrating preservation of mucosal architecture. Scale bars for H&E represent 500 mm (low power) and 100 mm (high power). **(I)** Immunohistochemical (IHC) analysis of diverticulitis-derived tissue at T0 and T48 hours. Markers include HIF-1a (metabolic/hypoxic adaptation), and KI-67 (cellular proliferation).

Most specimens had similar basal glucose consumption rates of approximately 0.1 mg/h/g. Two exceptions were observed: one diverticulitis-derived specimen (**Figure 4D**) showed a higher initial rate of approximately 0.3 mg/h/g, while one IBD-derived specimen (**Figure 4A**) showed a lower basal rate of approximately 0.035 mg/h/g. Lactate production followed a similar temporal pattern across all colon specimens.

In the IBD-derived small bowel specimen, the basal glucose consumption rate (∼0.08 mg/h/g) was comparable to that of diverticulitis-derived colon specimens at 4 hours. In most cases the glucose consumption rate increased over time, except for one specimen. Lactate production in the IBD and diverticulitis-derived specimens followed a similar temporal trend, depicting an increase in lactate production after the initial 4 hours and stabilizing at 48 hours.

Cumulative LDH released (normalized to tissue weight) showed an increasing trend indicating progressive tissue death or turnover as seen in **Figures 4C & 4F**. To understand how LDH might correlate with tissue death, we quantified the total LDH from diverse gut samples and created a standard curve to convert perfusate LDH to percentage tissue death. Based on this model, the cumulative LDH release corresponded to less than 0.4% tissue damage after 48 hours of perfusion. This low level of injury was representative of most specimens, with two outliers located at the minimum range of the cohort.

### Histological analysis of tissue

Preservation of tissue architecture and epithelial barrier integrity serves as the primary metric of successful perfusion. Representative H&E-stained sections from small bowel and colon specimens, acquired before and after perfusion, are shown in Figure 4G and 4H, respectively. Histological analysis demonstrated broad maintenance of intestinal architecture across multiple tissue layers, although a detectable progression of mucosal injury was observed after the 48-hour normothermic perfusion period. Prior to perfusion, the small bowel tissue exhibited well-preserved macro-architecture, where villus morphology and mucus production appeared to be intact (**Figure 4G**, left). Following perfusion, the most external layers of the lumen wall, such as muscularis propria, submucosa, and muscularis mucosa were well preserved, but with some signs of intercellular edema characterized by white spaces between cells within the aforementioned tissue compartments. The mucosa, especially the epithelial component, exhibited heterogeneity across the tissue section. While crypt architecture remained well-preserved in most of the tissue, clear morphological changes indicative of severe metabolic or inflammatory stress were observed in approximately one-third of the examined areas. This was characterized by mucosal sloughing, epithelial attenuation, denudation, disorganized crypt architecture, and fragmented lamina propria accompanied by significant submucosal edema (**Figure 4G**, right).

Similar to the small bowel IBD specimen, colonic tissue obtained from diverticulitis margins at T0 exhibited pristine architectural integrity (**Figure 4H**, left). This section featured tightly packed, parallel crypts and the presence of mucosa-associated lymphoid tissue (MALT, red arrow), corresponding to organized nodules of lymphocytes within the colonic mucosa. Following perfusion, overall preservation of the major tissue layers was observed, including the submucosa and muscularis propria, indicating effective vascular support. The epithelial layer and lamina propria of the mucosa were partially preserved, with the presence of packed, parallel crypts, like before perfusion. However, some other regions demonstrated increased interstitial edema, crypt distortion, and surface epithelial attenuation with significant surface epithelial lifting. Despite this regional heterogeneity, this specimen demonstrated superior overall resilience compared to the other specimens. Cell viability was further validated via Calcein AM/PI staining on post-perfusion biopsies, which confirmed predominantly viable cells (green) over dead (red) cells.

Next, we investigated the expression pattern of two key genes –HIF-1α to assess if tissue was hypoxic, and Ki-67 as a proliferation maker for intestinal stem cells. IHC analysis targeting HIF-1α and Ki-67 showed a reduction in both markers after perfusion (**Figure 4I**). Interestingly, we found that HIF-1α was highly expressed in stromal cells of the lamina propria before perfusion but was weaker and more diffuse post perfusion. One hypothesis to explain this observation is that the T0 sample was taken post ischemia which could have increased HIF expression, and perfusion may mitigate that effect. Unsurprisingly, Ki-67 had a strong stainingin glandular crypt epithelium and within the germinal center of the lymphoid follicle in the T0 sample, reduced expression in the T48 sample. This suggests that although perfusion preserved overall tissue viability, there was significant loss of intestinal stem cells over the 48 hours period.

### Real-time immunological profiling during NMP

Because the bowel is one of the most immunologically active organs, with constant immune cell trafficking, we sought to characterize how the immune cells within each specimen behaved on the NMP system. Thirteen cytokines were selected for their established roles in inflammation, IBD progression, and tissue repair. Perfusate cytokine levels were measured every 24 hours using a multiplexed bead-based immunoassay, which revealed time-dependent changes in secretion across most specimens (**Figure 5A–B**). While TNF-α, IL-1β, MCP-1, IL-18, and IL-8 were consistently detected in all samples, other cytokines appeared in only a subset. IL-6, IL-8, and MCP-1 reached the highest levels, particularly at later timepoints, likely reflecting innate immune cell activation (**Figure 5A–B**). Because this pattern was observed across all samples irrespective of disease state, the most probable cause is tissue damage arising from surgical trauma rather than disease-specific inflammation. In some samples we detected T cell–associated cytokines IL-2 and IL-17A, while IL-10 was present in three samples, possibly indicating compensatory immunosuppressive and tissue-repair activity. Most cytokines peaked at 24 hours and declined by 48 hours, whereas IL-1β and IL-6 showed a consistently increasing trend. These results are preliminary and cannot be used to interpret the immune pathways active in samples. Nevertheless, they demonstrate that the tissue remains immunologically active throughout the perfusion period.

**Figure 5.**
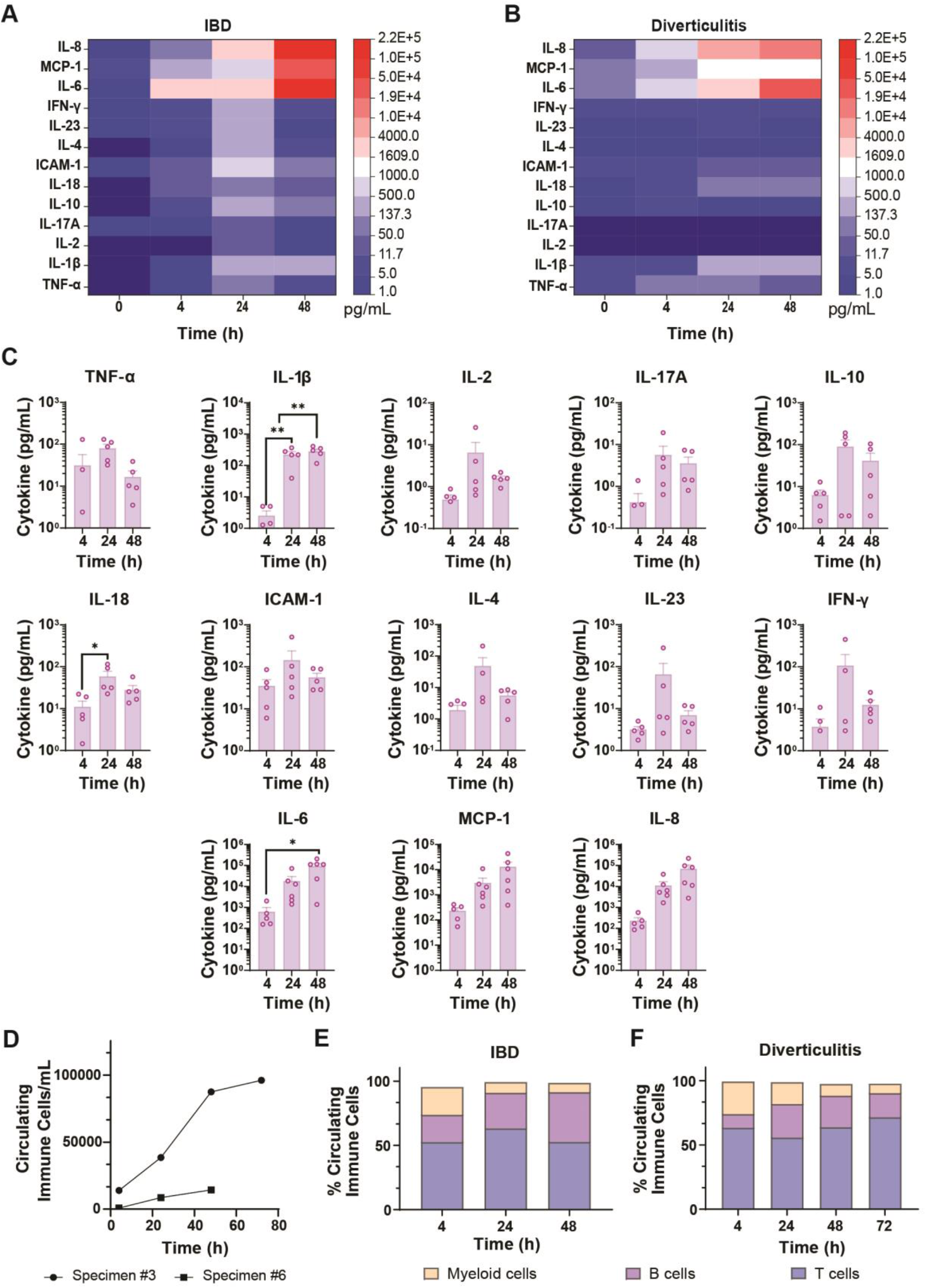
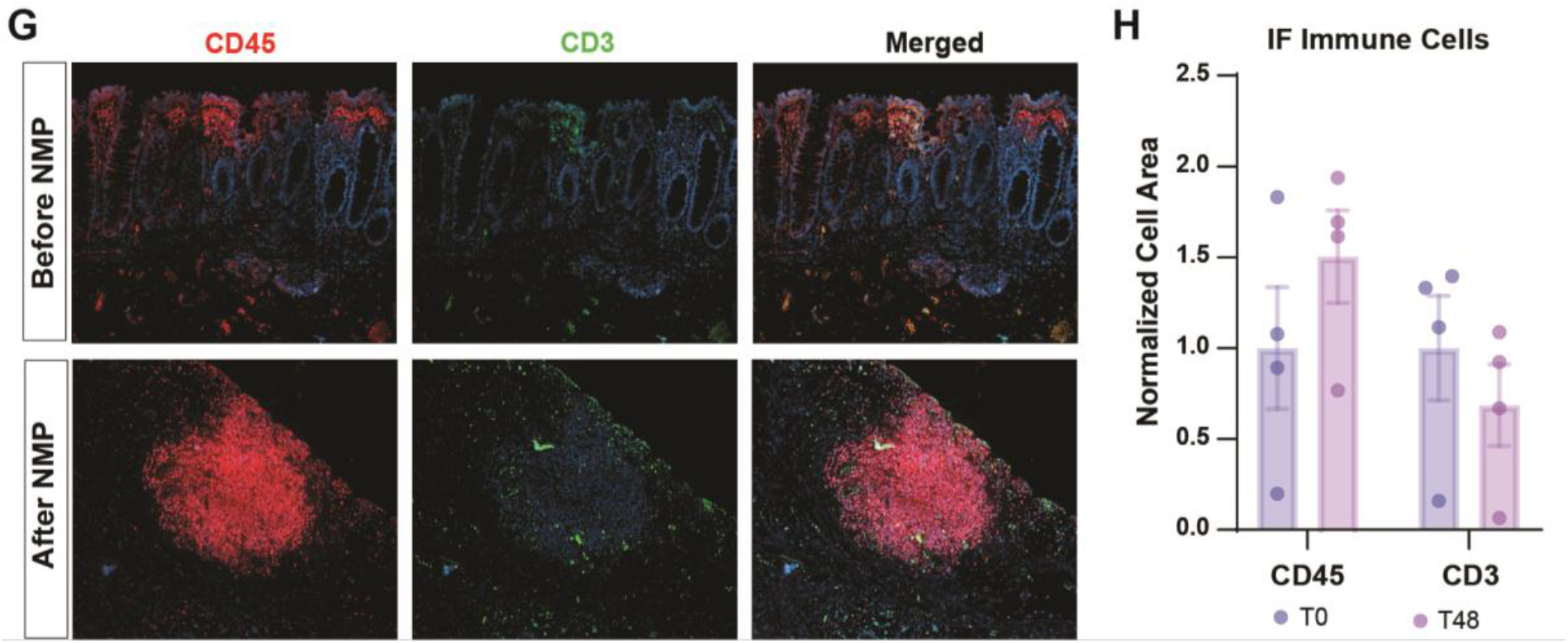
Pathology-specific immune signatures and secretome dynamics during NMP. **(A-B)** Heatmaps showing longitudinal circulating cytokine profiles of A) IBD and B) diverticulitis-derived specimens, illustrating distinct inflammatory trajectories. **(C)** Average of circulating cytokines per time point of all 5 experiments, except for IL-6, MCP1, and IL8, which represent the average of all 6 experiments. **(D)** Cumulative circulating immune cells of the perfusion of two specimens. Immune cells subset profiles demonstrating the three main leukocyte populations in E) the IBD and F) the diverticulitis-derived specimens. **(H)** Immunofluorescence (IF) of tissue sections at T0 and T48h (CD45: red; CD3: green; DAPI: blue), of) diverticulitis-derived specimen. **(I)** Quantification of immune cells infiltration in tissue, comparing before (T0) and after (T48) perfusion by normalizing cell area to T0. Data are presented as mean ± SEM. A One-Way ANOVA test was used for the statistical analysis of circulating cytokines; while Student’s t-test was applied to the histological immune cell quantification. Statistical significance was defined as p<0.05, and 95% confidence intervals are reported where indicated.

Since gut is a prime site for immune cell trafficking and surveillance, we expected resident immune cells within the tissue might extravasate into the perfusate during NMP. Accordingly, we analyzed circulating immune cell phenotyping across two representative specimens, one from IBD (sample #6) and the other from diverticulitis (sample #3). We note that this was not possible for all samples as we needed to include in-line cell filters in some samples where circulating cells clumps were noted to prevent embolism.

The immune cells from the perfusate were stained for further characterization using flow cytometry. The target molecules were the following: CD45, CD3, CD19, CD4, CD8a, CD11b, and CD66b. There was a pronounced increase over time in CD45+ leukocyte levels in the perfusate from the diverticulitis-derived specimen perfusion, with cumulative values reaching up to 96130 cells/mL at 72 hours of perfusion, while CD45+ leukocyte levels were comparable lower when measured from the IBD-derived specimen perfusion, reaching values up to 14370 cells/mL (**Figure 5D**). Representative leukocyte subset distribution is presented in **Figure 5E** and **5F** for circulating cells in perfusates obtained from perfusing IBD and diverticulitis-derived specimens. The dominating subset was CD45+CD3+ T cells in both cases, representing over 50% of the total immune cell population. CD45+CD19+ B cells subset showed an increase over time, and the opposite trend was observed in CD45+CD3-CD19-Myeloid cells in both cases. Histological examination of these specimens at baseline (T0) showed the presence of immune cells, CD45+ and CD3+ cells (**Figure 5G**). 5The quantification of the area of CD45+ cells and CD3+ cells was used to determine the shift in immune cell population in the tissue affected by NMP. The analysis of 4 different samples yielded an increase in CD45+ cells and a decrease in CD3+ cells when compared to tissue before perfusion. There was no statistical difference between before and after perfusion.

## DISCUSSION

The lack of high-fidelity experimental platforms that accurately recapitulate human intestinal physiology remains a primary bottleneck in the translation of therapies for inflammatory bowel disease (IBD) and diverticulitis.55,56 In this study, we established the feasibility of sustaining patient-derived human intestinal specimens within a custom NMP platform for 48-72 hours. By preserving the native mesenteric vasculature and integrated immune architecture (i.e. GALT), this "organ avatar" bridges the gap between simplistic in vitro systems and the translational shortcomings of animal models, thereby advancing intestinal NMP from a technical proof-of-concept toward a robust experimental system capable of capturing longitudinal biological events.

A major technical challenge in establishing ex vivo normothermic perfusion was the identification and cannulation of suitable arterial and venous vessels within the mesentery after tissue resection. This was particularly difficult in small bowel-derived specimens, in which the limited amount of attached mesentery and the small caliber of the vessels made reliable identification and cannulation of the vasculature challenging. On the other hand, some specimens contained substantial mesenteric adipose tissue, which obscured the vessels and complicated their dissection. Although both an artery and a vein could be identified in most specimens, surgical transection and tissue manipulation during resection and vessel preparation frequently resulted in multiple open vascular branches or areas of tissue damage that were difficult to seal completely. Fluoroscopic angiography was useful for identifying major sites of extravasation, and the application of PreveLeak surgical sealant reduced the extent of leakage; however, complete prevention of perfusate loss was not achievable. Persistent leakage likely contributed to the progressive reduction in venous return observed during perfusion.

A key accomplishment of organ perfusion was the multiscale preservation of different tissue levels across the intestine, including the serosa, muscularis externa, submucosa, and mucosa. Tissue architecture was broadly maintained post-perfusion; however, regional morphological heterogeneity was overserved within the epithelial layer, along with vascular frailty at the mesentery compartment, evidenced by focal crypt loss and fluoroscopic extravasation, respectively. These phenomena are likely caused by microembolism or regional vasoconstriction in the thin capillaries, limiting terminal oxygen delivery over time. Additionally, further optimization of perfusate composition may be required to support intestinal stem cells survival.

Interestingly, while bacterial contamination was detectable in the perfusate in samples #1 and #2, where optimized infection control was not implemented, there was no histological evidence of bacterial translocation into the tissue or overt infection-mediated structural damage. Instead, microbial overgrowth primarily impacted tissue viability indirectly through metabolic competition for glucose, suggesting that robust perfusion can mitigate the consequences of contamination if substrate delivery remains sufficient.

This platform could potentially capture patient-specific pathophysiology. Preliminary data demonstrated a distinct difference between cytokine profiles from different patients. Together with the implementation of personalized infection control strategies, these findings provide initial evidence that the platform preserves patient-to-patient variability, an important feature for its future development as a patient-specific drug-testing platform.

In summary, this validated NMP platform represents a significant advancement in preclinical experimental platforms by enabling metabolically active tissue investigation with greater fidelity than cold storage. The integration of epithelial, immune, and vascular compartments turns this platform into a holistic tool for personalized medicine and multidimensional therapeutic evaluation.

While this study establishes a foundation for long-term ex vivo modeling, several limitations remain. Our sample size (n = 6 for the finalized configuration) reflects the immense technical complexity of using fresh human surgical discards. While our iterative optimization (n = 11) informed the platform’s stability, larger cohorts are required to generalize these patient-specific responses. Furthermore, the inherent delay between organ procurement and the initiation of NMP introduces a period of cold ischemia. Given the intestine’s sensitivity to ischemia insult compared to solid organs, this transition remains a critical variable that may influence subsequent ex vivo recovery.

Technically, this system lacks an automated-feedback loop for addressing the flow control and mitigating vascular resistance. This is largely due to the anatomical challenges of venous return in fragmented mesenteric networks, which inhibits vascular resistance calculation. In this regard, it is also important to note that as this system operates as a semi-open vascular circuit rather than a strictly closed system, relying on passive collection of interstitial effluent from the specimen, circulating biomarkers should be interpreted with caution, as the extravasation of perfusate from mesentery can contribute to observed high number of circulating cells and inflammatory cytokines. Critically, our current focus on vascular-only perfusion does not address luminal near-physiological preservation. In the native intestine, the epithelium is supported by both basal blood flow and luminal factors. The absence of active luminal circulation may "starve" the nutrient-absorptive cells on the apical surface, potentially exacerbating the epithelial degradation observed in the histological analysis.

Furthermore, the current system lacks an active oxygenation component, with perfusate oxygen levels reaching only approximately 20%. Although the mucosal surface and villus tips are physiologically maintained in a relatively hypoxic state, the submucosal and mesenchymal layers of the healthy intestine are normally well oxygenated. These considerations underscore the need to incorporate an oxygenator into the perfusion circuit using a gas mixture of 95% O₂ and 5% CO₂. Insufficient oxygen delivery may explain why NMP attenuates, but does not fully normalize, HIF-1α expression, suggesting persistent tissue hypoxia and a corresponding compromise in tissue viability.

Future iterations of the “organ avatar” should adopt a dual-perfusion approach, managing the mesenteric vasculature and the intraluminal environment simultaneously. The incorporation of a portable pump for hypothermic perfusion system during transportation could further mitigate ischemia. Such advancements would allow for the simultaneous management of the microbiome and intraluminal pressure, potentially achieving the high-fidelity mucosal stabilization required for multi-day disease modeling and preclinical drug testing.

## ACKNOWLEDGEMENT

The authors acknowledge Jenny Pattengill at the Mayo Clinic Arizona Histology Core Laboratory for tissue processing and H&E staining. The authors also acknowledge Arizona State University Histology Core Laboratory for assistance in IHC staining. Authors would like to acknowledge funding from Mayo Clinic Investment for Extramural Grants Award (MEGA) to S.B., H.C. and J.B., Mayo Clinic Arizona Presidential Funds to S.B., H.C. and W.A.F., and Gift from The Opatrny Family Foundation to S.B. and H.C.

## AUTHOR CONTRIBUTIONS

Conceptualization, interpretation, and writing by P.M.U., S.R.C., J.B., H.C., and S.B. Methodology, investigation, and data analysis by P.M.U, S.R.C., and D.H. Resources provided by D.C, L.E., M.G.N, S.Y., N.R., T.A.L., H.H.A., F.H., N.N., and J.T.B. Pathology report by K.M. Supervision and funding acquisition by W.A.F., J.B., H.C., and S.B.

## Notes

### Competing Interest Statement

The authors have declared no competing interest.

